# Association of Macular Pigment Density with Plasma Macular Carotenoids levels, And lipids in Indian patients with early Age-Related Macular Degeneration

**DOI:** 10.1101/2024.02.23.581712

**Authors:** Senthilkumari Srinivasan, Anand Rajendren, Bala Panneerselvam, Mohammed Sithiq Uduman

**Affiliations:** Department of Ocular Pharmacology, Aravind Medical Research Foundation (AMRF), #1, Anna Nagar, Madurai, INDIA; Retina and Vitreous Services, Aravind Eye Hospital, #1, Anna Nagar, Madurai, INDIA; Department of Biostatistics, Aravind Eye Hospital, #1, Anna Nagar, Madurai, INDIA

**Keywords:** South Indian Eyes, healthy eyes, Eyes with early AMD, macular pigment optical density, plasma LZ and serum lipids

## Abstract

**Purpose:** The objective of the present study was to investigate the relationship of macular pigment optical density (MPOD) with plasma carotenoids [(L) and (Z)] and serum lipids in South Indian young healthy volunteers and patients with early AMD.

**Methods:** Two hundred and fourteen (N= 214) study participants (Healthy control group (N) = 178; Early AMD group (N) = 36) were enrolled after getting their written informed consent. The MPOD of the study participants was assessed using MPS II (Electron Technology, UK) after completing their routine ocular examination. Serum lipids were measured by the standard technique. Plasma levels of L, Z, lycopene and beta-carotene were estimated by HPLC with PDA detector.

**Statistical analysis used:** Correlations among variables in serum, plasma and the MPOD were established using Spearman’s rho correlation coefficient.

**Results:** The overall mean MPOD in healthy control group and early AMD group was found to be 0.47 ± 0.16 (N= 178; 317 eyes) and 0.35 ± 0.22 (N=36; 38 eyes) at 1° eccentricity respectively and was found to be significantly low as compared to healthy control group (p<0.001). A strong positive association was found between plasma L, Z and L+Z and MPOD. Serum HDL showed a strong negative association with MPOD and other lipids showed very weak association. MPOD was unaffected by BMI.

**Conclusions:** MPOD is positively associated with plasma L,Z and L+Z, adding further evidence that additional intake of L/Z may be beneficial in delaying the risk of AMD in our population

## Introduction

Macular pigment (MP) comprises of dietary lutein (L), Zeaxanthin (Z) and mesozeaxanthin (MZ) which is mainly located in the ganglion cell layers and inner plexiform layer of sensory retina where it shields the retina from UV light induced photo-oxidative damage(1). MP reflects the levels of L and Z within and outside of the fovea and its distribution decreases approximately at 5° to 8° eccentricity (2,3). The macular concentration of carotenoids is 1000 times greater than that of blood (4). Since, these carotenoids cannot be synthesized de novo, it has to be obtained through diet and the intensity of MP varies accordingly.

Multiple epidemiological studies have shown that low plasma L and Z concentrations or dietary intake are associated with low MP density and increased the risk of age-related macular degeneration (AMD) (5). Previous meta-analyses showed that the dietary intake of L and Z could lead to a 4% reduction in the risk of developing early AMD, as opposed to a 26% reduction for late AMD, indicating that LZ might be more effective in reducing the risk of progression from early AMD to late AMD (6). In another study, the supplementation of lutein at 10 or 20 mg/day significantly increased MPOD and contrast sensitivity. The stratified analyses of the same showed that the increase in MPOD to be faster and greater with higher dose and longer treatment (7).

Though the prevalence of AMD in India is comparable with the western countries, the knowledge of MP density and its modifiable risk factors in Indian population is not very well studied. Raman et al. (2012) attempted to obtain the normative data from healthy South Indian eyes and also studied the status of MP in wet AMD eyes and found significantly low MPOD vales as compared to control eyes (8). This finding further necessitates the importance of understanding the role of macular carotenoids in preservation of vision in both health and diseased Indian eyes. Therefore, in the present study, we attempted to investigate the relationship between plasma carotenoids, serum lipids and MPOD in young healthy Indian eyes and also in patients with early dry AMD. The results of the present study revealed that, the mean MPOD value was comparable with the previous study reported from South India and males were found to have significantly higher values as compared to females. In addition, this study found a strong positive correlation between MPOD and plasma L+Z. This observation holds true even after expressing the plasma L+Z levels in relation to serum lipids. The other carotenoids didn’t show any association with MPOD. The MPOD of early AMD group was found to be significantly low as compared to healthy control group (p<0.001). There was no significant association between MPOD and age in both groups. Serum HDL showed a strong negative association with MPOD and other lipids showed very weak association. MPOD was unaffected by BMI. These findings adding further evidence that additional intake of LZ may be beneficial in delaying the risk of AMD in our population.

## Subjects and Methods

### Study Subjects

Healthy volunteers Group: Volunteers with the age-group ranging from 20-81 years were recruited from our organization April 2015 to March 2017 after obtaining the written informed consent.

Study volunteers with no clinical evidence of ocular pathology, no dietary supplementation with LZ and visual acuity 20/40 or better were included in the study and those who were not willing to participate in the study; those with visually significant cataract and or macular disease on anterior & posterior segment photography and those who were on food supplementation rich in carotenoids were excluded from the study. Nine volunteers were excluded from the study due to their high fasting blood glucose (N=4), total serum cholesterol (N=2) and missing MPOD and Plasma LZ values (n=3).

#### Early AMD Group

Patients with early signs of AMD (drusen more than 63 μm) who were 49-79 years were included in this study. Patients with severe media opacity and any sign of retinopathy other than early AMD or shallow anterior chamber precluding mydriasis were excluded from the study.

The study protocol was approved by the standing Human Ethics Committee of our institution and was conducted in accordance with the Declaration of Helsinki. The written informed consents were obtained all study participants before their enrolment.

Demographic data including age, sex, education, and the anthropometric measurements such as height, weight, waist, and hip were collected for all study participants. All study participants underwent a detailed ophthalmic examination in the Retina Clinic of our hospital. Fundus colour photographs centred on macular area in each eye was documented before MPOD measurement. In addition, fasting blood samples were collected from study subjects to measure plasma carotenoids, serum lipids, blood glucose and haemoglobin. All samples were stored at -80° C till analysis. All the participants were asked to complete the questioners related to alcohol consumption, smoking status and the usage of any oral supplements.

### Macular Pigment Measurement

After a detailed ocular examination, all study subjects underwent MPOD measurements in both right and left eyes using macular pigment screener (MPS) II (Electron Technology, Cambridge, UK) as described in details elsewhere (9,10). Briefly, the MPOD test consists of two measurements: 1) central (0°) measurement and 2) peripheral (8°) measurement. The target consists of a 1° circular aperture in an integrating sphere which is uniformly illuminated with white light subtending approximately 30°. For central measurement, the study subjects were presented with a light stimulus of two alternating wavelengths (blue (465 nm) and green (530 nm)) and they were asked to report for the appearance of flicker (blue-green light) whereas peripheral measurement was achieved by fixating on a larger 1.75° red spot located at 8° horizontal eccentricity. The software of the MPS II automatically analyses the results upon response and gives three possible outcomes (accept, caution and reject). The readings of the subjects with unacceptable outcomes such as “caution” or “reject” were discarded and the study subject was asked to repeat it the following day until to get acceptable outcome (10). MPOD measurements of each study subject were performed by three trained investigators (SSK (Observer 1), AK (Observer 2) and BS). Forty eyes of twenty volunteers were selected to verify the repeatability (test-retest) of the instrument and to assess the inter-investigators (variability using Bland-Altman method before data collection (Bland and Altman, 1986). Readings with more than 0.2 standard deviations between eyes were excluded from the study.

#### Blood Measurements

Hemoglobin was estimated by cyan-methaemoglobin method using automated blood cell counter (Sysmex KX21, USA) and the other parameters were measured with a dry chemistry analyzer (Johnson & Johnson Vitros 250, Ortho-Clinical Diagnostics Inc., NY, USA).

Plasma lutein and zeaxanthin were estimated by high performance liquid chromatography (HPLC) method as described previously after extracting carotenoids using hexane. Sudan III was used as an internal standard (11). The plasma carotenoids were estimated using Shimadzu Prominence HPLC system with PDA detector (Shimadzu Corporation, Kyoto, Japan). Betacarotene and lycopene were estimated by the method as described by Kandar et al., (2013) with slight modifications (12). Briefly, the separation of betacarotene and lycopene was achieved with the mobile phase consisted of methanol and ethanol (75:25 v/v) pumped into Luna C8 (250X5 mm; 5μm) at the flow rate of 0.6 ml /minute. The column oven temperature was maintained at 40° C. Lutein, zeaxanthin, betacarotene, lycopene and Sudan III were identified by their absorption spectra and retention times using in-built library matching facility in the PDA detector.

### Statistical methods

All statistical analysis was done using statistical software STATA 11.0 (StataCorp, Texas, USA). Age, BMI and Plasma measurements are presented with Mean, standard deviation (SD) and Median with minimum and maximum. Distribution of the data was assessed for normality using Shapiro-Wilk test and Box-Whisker plot. Plasma measurements were compared between gender and BMI using Student’s t-test for normally distributed data and Mann-Whitney U test for skewed data. Correlations among variables in serum, plasma and the MPOD were established using Spearman rank order correlation (rho) or Pearson correlation coefficient (r) as appropriate. P<0.05 was considered statistically significant.

## Results

Of the 250 participants (Healthy volunteers group: 200; Early AMD Group: 50) enrolled, 36 participants (22 from healthy control group and 14 from early AMD group) were withdrawn from the study. Therefore, the data was collected from a total of 214 (Healthy control group (N) = 178; Early AMD group (N) = 36) participants. Of the remaining 178 participants in healthy control group, 72 (40.4%) were males and 106 (59.6%) were females with the age ranged from 20-81 years. Descriptive statistics of the study participants are given in supplementary table 1.

### Healthy control group

In healthy control group, a total of 317 (161 right eyes and 156 left eyes) eyes were available for MPOD measurement with a mean of 0.47 ±0.16. Measurement of MPOD between right eyes (0.48 ±0.16) and left eyes (0.46 ±0.16) did not differ significantly (p=0.356). There was a marginal increase from 20 – 30 years to 31-40 years followed by decline in MPOD but such decline was not significant (p=0.600; Figure 1, Table 1).

**Figure 1.**
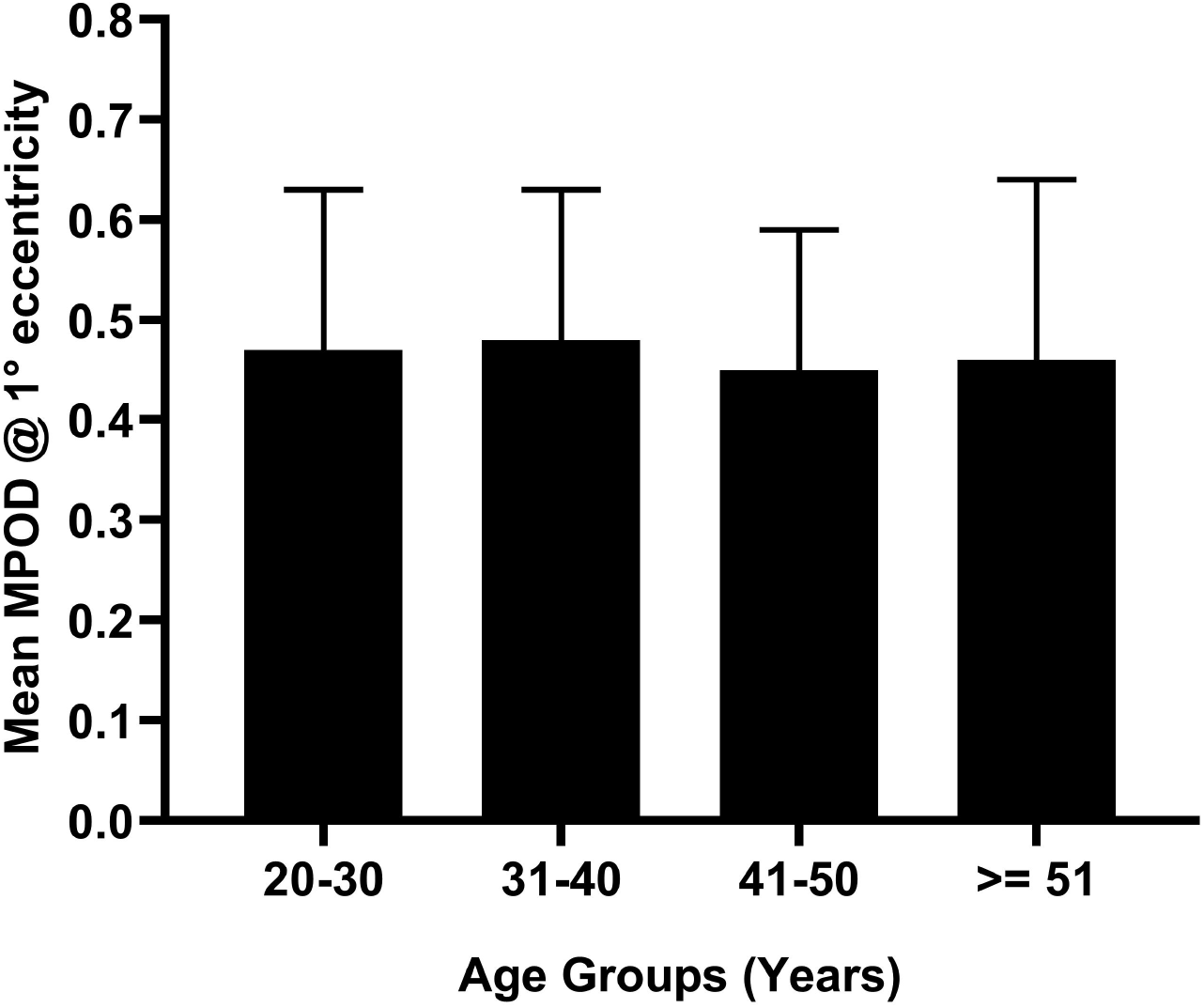
Mean MPOD at 1° eccentricity in 4 different age groups. The mean MPOD at 1° eccentricity for the different age groups of Healthy volunteers. There was a marginal increase from 20 – 30 years to 31-40 years followed by decline in MPOD but such decline was not significant (p=0.6; Supplementary Table 1).

Increasing trend of the biochemical indexes such as total cholesterol (p=0.005) and triglycerides (p=0.001) were observed with increase in age and were found to be statistically significant. However, HDL levels did not show any significant change with age. The plasma carotenoids such as L (p=0.003) and L+Z (p=0.012) showed increasing trend with increasing age and was found to be statistically significant. The plasma levels of other carotenoids such as Z, lycopene and beta-carotene in older groups (>= 51) were higher as compared to younger group (20-30 years).

MPOD Vs Gender: The mean MPOD of males and females with their biochemical and plasma data are summarized in Table 2. In this group, females were found to have low MPOD (0.44±0.14) as compared to males (0.52 ±0.17; p<0.001; Table 2).

Serum lipids and Plasma carotenoids based on gender: The serum total cholesterol, triglycerides (p=0.024) and LDL (p=0.042) were found to be high in males as compared to females and such difference was found to be statistically significant except total cholesterol (Table 2). Females were found to have higher plasma carotenoids as compared to males. The plasma betacarotene was found to be statistically significant (p=0.001).

### ***Correlation between MPOD and serum biochemical indexes, plasma carotenoids and*** Healthy volunteers

A Pearson correlation coefficient was used to describe the relation between MPOD, serum lipids, plasma carotenoids (L,Z, L+Z, lycopene and betacarotene) and BMI and the results are summarized in Table 3. MPOD was found to have strong positive correlation with plasma L,Z and L+Z(p<0.001; Table.5). The other carotenoids were found to have no correlation with MPOD. Serum HDL was found to have a strong negative association with MPOD (p<0.001). Similarly, serum total cholesterol, triglycerides were found to have negative correlation whereas LDL showed positive correlation with MPOD but these correlations were not statistically significant.

### Early AMD Group

In the early AMD group, the mean (SD) age was found to be 66.64 ±8.51 years with 28 (77.8%) were males and 8 (22.2%) were females. The patient characteristics are summarized in supplementary table 1. The mean MPOD of eyes with early AMD was found to be 0.35 ± 0.22 which was significantly lower as compared to healthy volunteer group (0.47 ±0.16; p<0.001; Supplementary Table 1; Figure 2).

**Figure 2:**
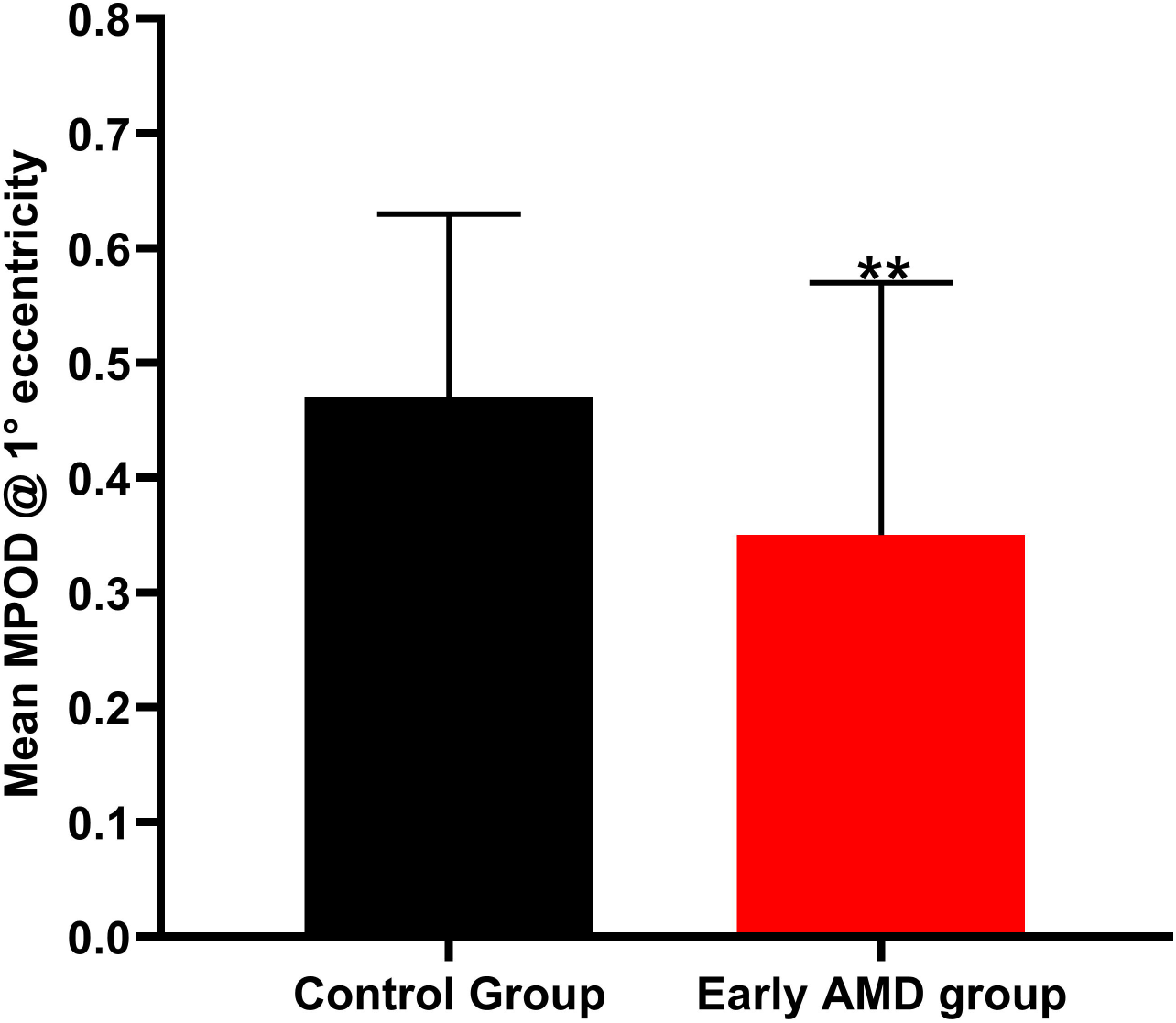
Comparison of Mean MPOD between Healthy control group and Early AMD Group. The mean MPOD of eyes with early AMD was found to be 0.35 ± 0.22 which was significantly lower as compared to healthy volunteer group (0.47 ±0.16; p<0.001; Supplementary Table 1). **p< 0.001; Student’s-t-test

The levels of serum triglycerides (p<0.001) and plasma L (p=0.002) values were significantly higher in early AMD group as compared to healthy volunteers group whereas the plasma levels of Z was significantly lower in AMD group (p=0.011; Supplementary Table 1). The correlation between MPOD, serum lipids, plasma carotenoids and BMI are summarized in Table 4. The MPOD was found to have positive correlation with plasma L and L+Z in early AMD group but such correlation was not statistically significant whereas lycopene showed a strong negative correlation (p<0.05). Serum lipids did not have any significant correlation with MPOD.

## Discussion

This is the first study to investigate the association of MPOD with plasma carotenoids and serum lipids in South Indian eyes (both healthy and early AMD eyes). Increasing evidences suggest that, insufficient macular pigments in human retina have been associated with retinal pathologies such as AMD. Since the prevalence of AMD in India is comparable with the western countries, the knowledge of MP and the factors responsible for its alteration is not well studied in our population. Therefore, in the present study, the relationship between normative data of MPOD and plasma L, Z and serum lipids was investigated.

In our present study, the MPOD values from all study participants were obtained using MPS II which is based on the principle of herteroflicker photometry. Several techniques have been used to estimate MPOD indirectly in living subjects such as herteroflicker photometry, raman spectroscopy, Fundus autofluorescence and reflectance photometry (13). Among such techniques, herteroflicker photometry which is a psychophysical technique claims to be the most accurate and reproducible (14,15). Repeatability was calculated for both instruments and the observer before the initiation of the study and found good agreement with the mean difference of -0.3 to 0.23 and -0.25 to 0.19 respectively at 95% confidence interval. This study found the overall mean MPOD at 1° eccentricity was 0.47 ± 0.15 which is comparable with the findings of Raman et al. (2012) in the same population (8). They reported the overall mean MPOD at 1° eccentricity was 0.37 ± 0.22 by considering all five age-groups (20-29,30-39,40-49,50-59 and >60 years). However, the reported mean MPOD was 0.51, 0.55 and 0.52 in the age groups of 20-29, 30-39 and 40-49 respectively. By taking the average of these three values, the MPOD was found to be 0.53. The negligible difference of 0.06 could be due to variation in the method of measurement and instrument used for the analysis. In another study, the mean MPOD measured using MPS II was 0.47± 0.1 with the age group of 20-34 years which is in good agreement with the observation of the present study (10). However, the ethnicity is different from the present study (Caucasians Vs Indians).

The results of MPOD on sex differences are very conflicting. In the present study, the males were found to have significantly higher MPOD as compared to females. This finding is in common with several other studies including the study reported in the same cohort (3,8,16). In contrast, males showed marginally lower plasma L+Z as compared to females but such difference was not statistically significant (Supplementary Table 1). It is possible that, the high MPOD could be attributed to the influence of testosterone on retinal uptake of carotenoids. Very interesting observation by Toomey et al (2015) suggest the involvement of testosterone in regulating the uptake, transport, and/or accumulation of carotenoids in the avian retina and driving seasonal patterns of accumulation (17). However, it would be interesting to study the correlation between testosterone levels and MPOD in males which would help us to understand the intricacies involved in carotenoid metabolism and retinal uptake based on gender.

The mean plasma L + Z in our present study was 0.49 μmol/L (0.18 – 1.00 μmol/L) which correlates well with the normative value reported by the Third National Health and Nutrition Examination Survey (NHANES III - 0.41 μmol/L) (18). The present study found a strong positive significant association between central MPOD and plasma L and L+Z (r = 0.149, r = 0.142; p<0.05 respectively). This is the first study which reported such association in Indian population. Plasma L+Z showed a very strong association with serum total cholesterol (r = 0.26; p<0.01) which is in good agreement with the previous studies (19,20). It is reported that, the primary transporter of xanthophylls such as L and Z in serum is HDL whereas carotenes are transported by LDL (21). The ratio of L in HDL to L in LDL is 3:1 indicating HDL lipoprotein is predominantly involved in the transport of L in serum (22). Since, there is an inverse association between serum triglycerides and serum HDL, this expected inverse association was found in the present study but it was not statistically significant. The factors which influence the levels of serum HDL will have an impact on plasma L and hence MPOD. Such inverse association was observed between serum triglycerides and MPOD. This suggest that, the individuals with high triglycerides are at the risk of low serum L, MPOD and likely to develop AMD. Further studies in this line are warranted to identify such link in our population.

It is interesting to note that lycopene levels showed a strong negative association with MPOD in patients with early AMD in the present study. There are some evidences suggest the role of lycopene in AMD (5,23,24). However, the protective role of lycopene in macular health is not well studied and future studies will be designed to address this hypothesis.

The low MPOD values in eyes with early AMD could be due to the small sample size. Despite of the repeated instructions about the test procedures, many elders enrolled in the study were not able to understand the test and hence they were not able to complete the test. This could also contribute to the poor association of MPOD to plasma carotenoids. Future studies with alternate measurement of MPOD such as fundus autofluorescence may help in this regard.

In conclusion, this is the first study to demonstrate a strong positive correlation between MPOD and plasma carotenoids (L+Z) in our healthy Indian population and patients with early AMD. The estimated MPOD was comparable with the study reported earlier in the same population. MPOD values of healthy groups was not affected by age, however, females were found to have significantly low MPOD values as compared to males which in contrary to the previous study. Plasma L, Z and L+Z showed a strong positive association with MPOD and such association was not reported earlier in our population whereas serum HDL showed a strong negative association with MPOD. In case of eyes with early AMD, we found a weak association between plasma carotenoids and MPOD, however lycopene showed a strong negative association. The low MPOD values in patients with early AMD emphasize the importance of additional intake of L/Z may be beneficial in delaying the risk of AMD in our population.

## Author contributions

Conceptualization: Senthilkumari Srinivasan; Methodology: Senthilkumari Srinivasan, Anand Rajendren, Bala Panneerselvam; Formal analysis and investigation: Senthilkumari Srinivasan, Mohammed Sithiq Uduman; Writing - original draft preparation: Senthilkumari Srinivasan, Mohammed Sithiq Uduman; Writing - review and editing: Senthilkumari Srinivasan, Anand Rajendren, Mohammed Sithiq Uduman; Funding acquisition: Senthilkumari Srinivasan; Resources: Senthilkumari Srinivasan, Anand Rajendren; Supervision: Senthilkumari Srinivasan.

## Acknowledgement

The authors acknowledge the study participants for their kind co-operation in this study. The study was supported by Indian Council of Medical Research, New Delhi, INIDA (No.5/4/6/3/Oph/2013-NCD-II) and the equipment cost was supported by Aravind Eye Foundation (AEF), New York.

