## Supplementary material for "Association of Macular Pigment Density with Plasma Macular Carotenoids levels, And lipids in Indian patients with early Age-Related Macular Degeneration": Table1, Table2, Table3, Table4

**Table 1:** Plasma Concentrations of carotenoids, biochemical indexes and MPOD in healthy controls of different age groups

| **Variables** | **Age group** | | | | **Total** | **P value ^a^** |
| --- | --- | --- | --- | --- | --- | --- |
|  | **20 – 30 years** | **31 – 40 years** | **41 – 50 years** | **Above 50 years** |  |  |
| **BMI**  n  Mean(SD)  Median  Min-Max | 65  22.87(2.88)  23.14  15.60 – 31.35 | 38  26.08(3.54)  26.00  18.70 – 33.90 | 15  27.24(5.58)  27.41  18.50 – 39.26 | 16  26.12(4.02)  25.30  20.10 – 37.34 | 134  24.66(3.97)  24.66  15.60 – 39.26 | **<0.001** |
| **MPOD**  n(**eye-wise**)  Mean(SD)  Median  Min-Max | 141  0.47(0.16)  0.46  0.12 – 0.98 | 90  0.48(0.15)  0.46  0.12 – 0.84 | 45  0.45(0.14)  0.41  0.22 – 0.79 | 41  0.46(0.18)  0.46  0.02 – 0.89 | 317  0.47(0.16)  0.46  0.02 – 0.98 | 0.600 |
| **Total cholesterol**  n  Mean(SD)  Median  Min-Max | 76  156.63(23.75)  153.00  111.00 – 219.00 | 44  157.11(33.16)  154.50  21.00 – 214.00 | 22  169.05(20.23)  164.00  132.00 – 205.00 | 24  176.83(27.24)  183.00  120.00 – 222.00 | 166  161.32(27.50)  158.00  21.00 – 222.00 | **0.005** |
| **HDL**  n  Mean(SD)  Median  Min-Max | 75  43.04(6.17)  43.00  23.00 – 58.00 | 40  43.53(15.11)  41.00  25.00 – 125.00 | 21  44.29(4.81)  45.00  35.00 – 53.00 | 24  42.75(6.83)  41.50  31.00 – 59.00 | 160  43.28(9.14)  42.50  23.00 – 125.00 | 0.938 |
| **Triglycerides**  n  Mean(SD)  Median  Min-Max | 75  93.92(39.63)  81.00  14.00 – 265.00 | 40  115.65(45.36)  99.00  51.00 – 239.00 | 21  127.62(62.06)  110.00  54.00 – 255.00 | 24  130.75(52.80)  124.00  69.00 – 288.00 | 160  109.30(48.56)  94.00  14.00 – 288.00 | **0.0009** |
| **LDL**  n  Mean(SD)  Median  Min-Max | 75  95.27(21.15)  91.00  44.00 – 141.00 | 40  94.5(23.63)  94.00  40.00 – 134.00 | 21  96.38(19.12)  93.00  40.00 – 130.00 | 24  108.29(25.07)  112.50  61.00 – 145.00 | 160  97.17(22.46)  93.50  40.00 – 145.00 | 0.070 |
| **Plasma L**  n  Mean(SD)  Median  Min-Max | 75  155.68(58.56)  143.15  52.60 – 373.91 | 36  163.55(52.64)  166.93  65.56 – 341.95 | 20  191.10(66.35)  178.18  96.57 – 322.17 | 22  205.67(76.01)  185.31  110.06 – 392.24 | 153  169.35(63.32)  154.27  52.60 – 392.24 | **0.003** |
| **Plasma Z**  n  Mean(SD)  Median  Min-Max | 75  117.11(48.08)  109.16  49.89 – 259.66 | 36  113.38(47.99)  110.13  34.33 – 244.36 | 20  133.66(59.25)  121.04  59.59 – 287.11 | 22  137.17(60.46)  125.40  72.39 – 336.15 | 153  121.28(51.76)  115.47  34.30 – 336.15 | 0.209 |
| **Plasma L + Z**  n  Mean(SD)  Median  Min-Max | 75  272.78(92.26)  258.08  104.14 – 567.17 | 36  276.93(86.80)  286.27  124.08 – 466.16 | 20  324.76(116.46)  313.21  166.65 – 571.4 | 22  342.84(129.61)  318.46  186.89 – 728.39 | 153  290.63(103.17)  268.23  104.10 – 728.40 | **0.012** |
| **Plasma Lycopene**  n  Mean(SD)  Median  Min-Max | 65  354.14(306.38)  234.27  4.21 – 975.92 | 31  303.81(266.35)  239.68  9.70 – 829.05 | 18  233.22(158.43)  219.40  13.61 – 496.59 | 18  414.99(263.22)  377.60  54.35 – 838.64 | 132  334.13(277.61)  253.67  4.21 – 975.92 | 0.318 ^b^ |
| **Plasma Betacarotene**  n  Mean(SD)  Median  Min-Max | 74  172.50(145.74)  115.63  10.10 – 606.1 | 35  147.14(110.28)  111.86  7.14 – 388.08 | 20  175.75(141.95)  184.62  17.06 – 486.75 | 20  245.54(154.79)  231.55  44.04 – 553.63 | 149  176.78(140.67)  140.93  7.14 – 606.10 | 0.128 ^b^ |
| **SD**-standard deviation;  ^a^ compared using ANOVA test between the age groups;  ^b^ compared using Kruskal-Wallis test between the age groups;  P-value less than 0.05 considered statistically significant (bold faced) | | | | | | |

**Table 2:** Gender wise distribution of Plasma Concentrations of carotenoids, biochemical indexes and MPOD in healthy controls

| **Variables** | **Gender** | | **P value** ^a^ |
| --- | --- | --- | --- |
|  | **Male** | **Female** |  |
| **BMI**  n  Mean(SD)  Median(Min-Max) | 52  25.13(4.07)  24.37(17.93 – 39.26) | 82  24.36(3.90)  24.85(15.60 – 37.11) | 0.272 |
| **MPOD**  n(**eye-wise**)  Mean(SD)  Median(Min-Max) | 128  0.52(0.17)  0.5(0.02 – 0.98) | 189  0.44(0.14)  0.41(0.12 – 0.84) | **<0.001** |
| **Total cholesterol**  n  Mean(SD)  Median(Min-Max) | 64  166.48(27.60)  163(119 – 219) | 102  158.09(27.08)  156.5(21 – 222) | 0.055 |
| **HDL**  n  Mean(SD)  Median(Min-Max) | 63  41.14(6.33)  42(23 – 53) | 97  44.67(10.37)  44(31 – 125) | **0.017** |
| **Triglycerides**  n  Mean(SD)  Median(Min-Max) | 63  120.05(55.12)  99(14 – 265) | 97  102.32(42.65)  87(52 – 288) | **0.024** |
| **LDL**  n  Mean(SD)  Median(Min-Max) | 63  101.65(23.47)  100(44 – 141) | 97  94.27(21.40)  93(40 – 145) | **0.042** |
| **Plasma L**  n  Mean(SD)  Median(Min-Max) | 58  162.22(61.43)  146.1(88.25 – 392.24) | 95  173.70(64.38)  166.59(52.6 – 351.17) | 0.278 |
| **Plasma Z**  n  Mean(SD)  Median(Min-Max) | 58  120.92(54.84)  109.99(34.33 – 336.15) | 95  121.49(50.09)  119.45(49.89 – 287.11) | 0.947 |
| **Plasma L + Z**  n  Mean(SD)  Median(Min-Max) | 58  283.14(106.89)  254.32(156.4 – 728.39) | 95  295.20(101.14)  270.43(104.14 – 571.4) | 0.485 |
| **Plasma Lycopene**  n  Mean(SD)  Median(Min-Max) | 49  315.17(257.88)  251.27(4.21 – 835.6) | 83  345.32(289.57)  277.49(13.61 – 975.92) | 0.536 ^b^ |
| **Plasma Betacarotene**  n  Mean(SD)  Median(Min-Max) | 57  129.36(116.48)  82.43(11.77 – 606.1) | 92  206.16(146.81)  195.11(7.14 – 576.81) | **0.001** ^b^ |
| **SD**-standard deviation;  ^a^ compared using Student’s t-test between the gender;  ^b^ compared using Mann-Whitney U test between the gender;  P-value less than 0.05 considered statistically significant (bold faced) | | | |

**Table 3:** The relation between MPOD, Serum Lipids, plasma carotenoids and BMI in Healthy volunteers

| **Variables** | **BMI** | **MPOD** | **Tchol** | **Triglycerides** | **HDL** | **LDL** | **L** | **Z** | **L+Z** | **Lycopene** |
| --- | --- | --- | --- | --- | --- | --- | --- | --- | --- | --- |
| BMI | 1.000 |  |  |  |  |  |  |  |  |  |
| MPOD | -0.021 | 1.000 |  |  |  |  |  |  |  |  |
| Tchol | 0.122* | -0.034 | 1.000 |  |  |  |  |  |  |  |
| Triglycerides | **0.226*** | -0.074 | **0.386**** | 1.000 |  |  |  |  |  |  |
| HDL | -0.062 | -0.161** | -0.196** | -0.106 | 1.000 |  |  |  |  |  |
| LDL | 0.043 | 0.024 | **0.867**** | 0.128* | -0.172** | 1.000 |  |  |  |  |
| L | 0.062 | 0.166** | **0.218**** | 0.093 | **-0.002*** | 0.158** | 1.000 |  |  |  |
| Z | -0.018 | 0.123* | **0.298**** | 0.007 | 0.094 | **0.221**** | **0.603**** | 1.000 |  |  |
| L+Z | 0.031 | 0.163** | **0.283**** | 0.062 | 0.045 | **0.207**** | **0.916**** | **0.872**** | 1.000 |  |
| Lycopene ^†^ | -**0.226**** | -0.026 | -0.097 | -0.075 | **0.191**** | -0.138* | **0.372**** | **0.361**** | **0.390**** | 1.000 |
| Beta-carotene ^†^ | -0.169** | 0.003 | 0.014 | -0.113 | **0.276**** | -0.037 | **0.516**** | **0.418**** | **0.513**** | **0.729**** |
| *significant (<0.05), **significant (<0.01);  Pearson correlation coefficient (r) and ^†^ Spearman rank order correlation coefficient values were reported | | | | | | | | | | |

**Table 4:** The relation between MPOD, Serum Lipids, plasma carotenoids and BMI in Early AMD group

| **Variables** | **BMI** | **MPOD** | **Tchol** | **Triglycerides** | **HDL** | **LDL** | **L** | **Z** | **L+Z** | **Lycopene** |
| --- | --- | --- | --- | --- | --- | --- | --- | --- | --- | --- |
| BMI | 1.000 |  |  |  |  |  |  |  |  |  |
| MPOD | 0.089 | 1.000 |  |  |  |  |  |  |  |  |
| Tchol | 0.044 | 0.109 | 1.000 |  |  |  |  |  |  |  |
| Triglycerides | 0.004 | 0.084 | 0.240 | 1.000 |  |  |  |  |  |  |
| HDL | 0.149 | 0.073 | 0.524 | -**0.449**** | 1.000 |  |  |  |  |  |
| LDL | -0.0001 | 0.061 | 0.845 | -**0.277*** | **0.613**** | 1.000 |  |  |  |  |
| L | 0.067 | 0.156 | 0.015 | -**0.340*** | **0.309*** | 0.081 | 1.000 |  |  |  |
| Z | -0.072 | 0.070 | -0.149 | -0.037 | -0.189 | -0.104 | 0.198 | 1.000 |  |  |
| L+Z | 0.041 | 0.160 | -0.027 | -**0.322*** | 0.232 | 0.046 | **0.963**** | **0.454**** | 1.000 |  |
| Lycopene ^†^ | -0.149 | -**0.436*** | -0.050 | -0.042 | -0.050 | -0.010 | 0.150 | **0.336*** | 0.177 | 1.000 |
| Beta-carotene ^†^ | -0.085 | -0.148 | 0.157 | -0.261 | **0.321*** | 0.260 | **0.335*** | **0.274*** | **0.334*** | **0.527**** |
| *significant (<0.05), **significant (<0.01);  Pearson correlation coefficient (r) and ^†^Spearman rank order correlation coefficient values were reported | | | | | | | | | | |
