## Supplementary Table for "Association of Macular Pigment Density with Plasma Macular Carotenoids levels, And lipids in Indian patients with early Age-Related Macular Degeneration"

**Supplementary Table 1:** Descriptive Statistics of Study Participants

| **Variables** | **Healthy control group** | | | **Early AMD group** | | | **P-value ^a^** |
| --- | --- | --- | --- | --- | --- | --- | --- |
|  | **n** | **Mean(SD)** | **Median**  **(Min - Max)** | **n** | **Mean(SD)** | **Median**  **(Min - Max)** |  |
| Age (years) | 178 | 35.76(12.73) | 33  (20 - 81) | 36 | 66.64(8.51) | 67  (46 - 79) | **<0.001** |
| BMI | 134 | 24.66(3.97) | 24.67  (15.6 – 39.3) | 31 | 25.06(4.49) | 25.23  (16.34 – 38.13) | 0.623 |
| MPOD | 317 | 0.47(0.16) | 0.46  (0.02 – 0.98) | 38 | 0.35(0.22) | 0.31  (0.02 – 0.89) | **<0.001** |
| Total Cholesterol | 166 | 161.33(27.50) | 158  (21 – 222) | 30 | 167.2(31.29) | 162.5  (106 – 234) | 0.293 |
| HDL | 160 | 43.28(9.14) | 42.5  (23 – 125) | 30 | 46.7(8.63) | 45  (31 – 61) | 0.060 |
| Triglycerides | 160 | 109.3(48.57) | 94  (14 – 288) | 30 | 154.4(93.22) | 129  (60 – 525) | **0.0001** |
| LDL | 160 | 97.18(22.46) | 93.5  (40 – 145) | 30 | 89.67(26.21) | 86  (40 – 135) | 0.104 |
| Plasma L | 153 | 169.35(63.32) | 154.27  (52.6 – 392.24) | 28 | 219.24(125.13) | 169.04  (98.85 – 623.08) | **0.002** |
| Plasma Z | 153 | 121.28(51.76) | 115.47 (34.33 – 336.15) | 28 | 94.93(37.76) | 85.75  (50.30 – 202.84) | **0.011** |
| Plasma L + Z | 153 | 290.63(103.17) | 268.23  (104.14 – 728.39) | 28 | 314.16(137.68) | 254.22  (149.67 – 735) | 0.295 |
| Plasma Lycopene | 132 | 334.13(277.61) | 253.67  (4.21 – 975.92) | 26 | 280.13(187.53) | 208.90  (48.21 – 824.70) | 0.918 **^b^** |
| Plasma Betacarotene | 149 | 176.78(140.68) | 140.93  (7.14 – 606.1) | 26 | 266.69(236.99) | 180.10  (12.74 – 889.16) | 0.075 **^b^** |
| **SD**-standard deviation;  ^a^ compared using Student’s t-test between healthy control and early AMD group;  ^b^ compared using Mann-Whitney U test between healthy control and early AMD group;  P-value less than 0.05 considered statistically significant (bold faced) | | | | | | | |
